# Global distribution of the rooting zone water storage capacity reflects plant adaptation to the environment

**DOI:** 10.1101/2021.09.17.460332

**Authors:** Benjamin D. Stocker, Shersingh Joseph Tumber-Dávila, Alexandra G. Konings, Martha B. Anderson, Christopher Hain, Robert B. Jackson

## Abstract

The rooting zone water storage capacity (*S*_0_) extends from the soil surface to the weathered bedrock (the Critical Zone) and determines land-atmosphere exchange during dry periods. Despite its importance to land-surface modeling, variations of *S*_0_ across space are largely unknown as they cannot be observed directly. We developed a method to diagnose global variations of *S*_0_ from the relationship between vegetation activity (measured by sun-induced fluorescence and by the evaporative fraction) and the cumulative water deficit (CWD). We then show that spatial variations in *S*_0_ can be predicted from the assumption that plants are adapted to sustain CWD extremes occurring with a return period that is related to the life form of dominant plants and the large-scale topographical setting. Predicted biome-level *S*_0_ distributions, translated to an apparent rooting depth (*z*_*r*_) by accounting for soil texture, are consistent with observations from a comprehensive *z*_*r*_ dataset. Large spatial variations in *S*_0_ across the globe reflect adaptation of *z*_*r*_ to the hydroclimate and topography and implies large heterogeneity in the sensitivity of vegetation activity to drought. The magnitude of *S*_0_ inferred for most of the Earth’s vegetated regions and particularly for those with a large seasonality in their hydroclimate indicates an important role for plant access to water stored at depth - beyond the soil layers commonly considered in land-surface models.

## Introduction

Water availability to vegetation exerts a strong control on the terrestrial carbon cycle [2, 30], surface-atmosphere exchanges [60], and the development of heat waves [46]. Under dry conditions and increasing cumulative water deficits (CWD, defined as the difference between evapotranspiration, ET, and precipitation, *P*, summed over continuous dry periods), water availability to vegetation becomes increasingly limiting, leading to reduced CO_2_ assimilation and transpiration [10], to accelerated heating of near-surface air [68], and - in extreme cases - to premature defoliation of water-stressed vegetation [5] and enhanced tree mortality [9, 25].

To sustain activity during dry periods and resist impacts of droughts, plants rely on water stored below the surface [26, 44]. The larger the rooting zone water storage capacity (*S*_0_), the longer plants can withstand soil moisture limitation during dry periods [66] and the lower their sensitivity to an increasing CWD. *S*_0_ is therefore a key factor determining drought impacts, land-atmosphere exchanges, and runoff regimes, particularly in climates with seasonal asynchrony in radiation and precipitation [23, 26, 44]. In models, *S*_0_ is commonly conceived as a function of the soil texture and the plants’ rooting depth (*z*_*r*_), limited to the soil depth [44, 59]. Recent research has revealed a substantial component of *S*_0_ and contributions to ET by water stored beneath the soil [48], in weathered and fractured bedrock [12, 43, 53] or groundwater [18, 28, 67]. Plant access to such deep moisture plays an important role in controlling near-surface climate [42, 57, 65], runoff regimes [26], global patterns of vegetation cover [37], and mitigating impacts of droughts [17].

However, *S*_0_ is impossible to observe directly across large scales and its spatial variations are poorly understood [39]. Gravimetric remote sensing provides information of terrestrial water storage variations but only at ~100 km spatial resolution [30]. Information from microwave remote sensing is limited to the top few centimeters of the soil, and derived root zone soil moisture estimates rely on extrapolations and assumptions for *z*_*r*_ [13]. Global compilations of local *z*_*r*_ measurements [7, 55] have resolved this observational challenge only partly because of their limited size (owing to prohibitively laborious data collection efforts), the importance of the (often unknown) groundwater table depth (WTD) in relation to *z*_*r*_ [18], and large documented variations in *z*_*r*_ across multiple scales. *z*_*r*_ varies across large-scale climatic gradients and the Earth’s biomes [7, 55], along topographical gradients and catchment-scale variations of the WTD [18], between individuals growing under different conditions [70], and among different growth forms and species growing in close vicinity [8, 55]. Empirical approaches for estimating the global *z*_*r*_ distribution made use of relationships between in-situ observations and climatic factors [54]. Modelling approaches for predicting *z*_*r*_ have conceived their spatial variations as the result of eco-evolutionary pressure and plants’ adaptation to the prevailing hydro-climate [21, 34, 58, 62, 74], or as being adapted to just buffer water demand by (observed) evapotranspiration and precipitation during dry periods [23, 69]. Such *mass-balance approaches* make use of the seasonality in radiation and precipitation, their relative timing, and the ensuing cumulative water deficit (CWD) as important driving factors for plants’ genetic or phenotypic adaptation of *z*_*r*_ and hence *S*_0_. However, their predictions remain untested against local *z*_*r*_ observations.

Despite its crucial role in controlling water and carbon fluxes and the scarcity of observations, practically all land surface models, climate models, terrestrial biosphere models, numerical weather prediction models, or hydrological models, and several widely used remote sensing-based models of vegetation productivity and evapotranspiration rely on a specification of *S*_0_ either directly as the depth of a “water bucket” (a simple model for water storage along the rooting zone), or indirectly through a prescribed rooting depth *z*_*r*_ and soil texture across the profile, or the shape of an empirical water stress function. Typically, water stored along the entire Critical Zone - the Earth’s outermost layer of biota and rocks that interacts with the atmosphere, the hydrosphere, and biogeochemical cycles - is not fully represented in models [28, 53]. As a consequence, spatial variations in *z*_*r*_ and *S*_0_ arise in models predominantly from prescribed variations in soil texture [38], from distinct parameter choices of *z*_*r*_ for different plant functional types (PFTs), and from the PFTs’ spatial distribution [14]. The evident plasticity of *z*_*r*_ and *S*_0_ along climatic and topographic gradients is typically ignored. Implications of this simplification may be substantial for the simulation of land-atmosphere coupling and drought impacts [28, 42, 57].

Here, we present a method for diagnosing *S*_0_ from the relationship between vegetation activity (measured by sun-induced fluorescence and by the evaporative fraction) and CWD. By fusing multiple time series of Earth observation data streams with global coverage, we estimate the global distribution of *S*_0_ at a resolution of 0.05°(~5 km). Using a *mass-balance approach* [23, 69] and field observations of *z*_*r*_ from a globally distributed dataset, we then show that the sensitivity of vegetation activity to water stress is strongly correlated with the magnitude of local CWD extremes, and that this reflects adaptation of the rooting depth to the local hydroclimate and water table depth.

## Methods

### Cumulative water deficit estimation

The cumulative water deficit (CWD) is determined here from the cumulative difference of actual evapotranspiration (ET) and the liquid water infiltration to the soil (*P*_in_). ET is based on thermal infrared remote sensing, provided by the global ALEXI data product at daily and 0.05° resolution, covering years 2003-2018. Values in energy units of the latent heat flux are converted to mass units accounting for the temperature and air pressure dependence of the latent heat of vaporization following [11]. *P*_in_ is based on daily reanalysis data of precipitation in the form of rain and snow from WATCH-WFDEI [71]. A simple snow accumulation and melt model [50] is applied to account for the effect of snowpack as a temporary water storage that supplies *P*_in_ during spring and early summer. Snow melt is assumed to occur above 1°C and with a rate of 1 mm d^−1^ (°C)^−1^. The CWD is derived by applying a running sum of (ET - *P*_in_) for days where (ET - *P*_in_) is positive (net water loss from the soil), and terminating the summation after rain has reduced the running sum to zero. All precipitation and snowmelt (*P*_in_) is assumed to contribute to reducing the CWD. This implicitly assumes that no runoff occurs while the CWD is above zero. The period between the start and end of accumulation is referred to as a CWD event. Within each event, data are removed after rain has reduced the CWD to below 90% of its maximum value within the same event. This concerns the analysis of SIF and EF (see below) and avoids effects of relieved water stress by re-wetting topsoil layers before the CWD is fully compensated. The algorithm to determine daily CWD values and events is implemented by the R package *cwd* [63].

### Estimating *S*_0_ from SIF and EF

As the ecosystem-level CWD increases, both gross primary production (GPP, ecosystem-level photosynthesis) and evapotranspiration (ET) are limited by the availability of plant access to water. Below, we refer to GPP and ET as a generic “vegetation activity” variable *X*(*t*). This principle can be formulated, in its simplest form, as a model of *X*(*t*) being a linear function of the remaining water stored along the rooting zone *S*(*t*), expressed as a fraction of the total rooting zone water storage capacity *S*_0_ [40]:

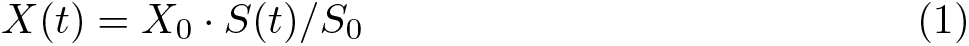

Following Eq. 1, *S*_0_ can be interpreted as the total rooting zone water storage capacity, or the depth of a “water bucket” that supplies moisture for evapotranspiration. Following [66] and with *X*(*t*) representing ET, the temporal dynamics during rain-free periods (where runoff can be neglected), are described by the differential equation

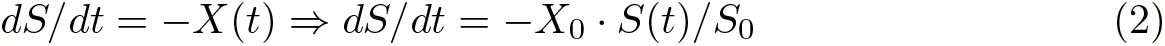

and solved by an exponential function with a characteristic decay time scale λ:

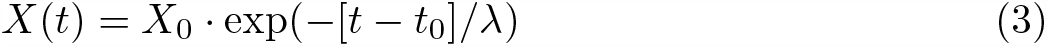

λ is related to *S*_0_ as *S*_0_ = λX_0_, where *X*_0_ is the initial ET at *S*(*t*_0_) = *S*_0_. In other words, the apparent observed exponential ET decay time scale λ, together with *X*_0_, reflects the total rooting zone water storage capacity *S*_0_.

Fitting exponentials from observational data is subject to assumptions regarding stomatal responses to declines in *S*(*t*) and is relatively sensitive to data scatter. Hence, resulting estimates of *S*_0_ may not be robust. With CWD= *S*_0_ − *S*(*t*) and Eq. 1, the relationship of *X*(*t*) and CWD(*t*) can be expressed as a linear function

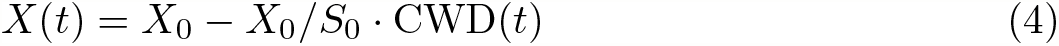

and observational data for *X*(*t*) can be used to fit a linear regression model. Its intercept *a* and slope *b* can then be used as an alternative, and potentially more robust estimate for *S*_0_:

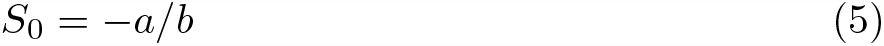

This has the further advantage that estimates for *S*_0_ can be derived using any observable quantity of vegetation activity *X*(*t*) (not just ET as in [66]) under the assumption that activity attains zero at the point when the CWD reaches the total rooting zone water storage capacity; i.e., *X*(*t*\*) = 0 for CWD(*t*\*) = *S*_0_.

Here, we use sun-induced fluorescence (SIF, [16]), normalised by incident shortwave radiation (WATCH-WFDEI data [71]), and the evaporative fraction (EF), defined as the ratio of evapotranspiration (ALEXI-ET data [27]) over net radiation (GLASS data [32]), as two alternative proxies for water-constrained vegetation activity *X*. ET and net radiation data are both provided at 0.05° and daily resolution.

*S*_dSIF_ and *S*_dEF_ were derived based on the linear relationship of SIF and EF versus CWD by applying Eq. 5. The relationship was derived for each pixel with pooled data belonging to the single largest CWD event of each year, and using the 90% quantile of EF and normalised SIF within 50 evenly spaced bins along the CWD axis. Binning and considering percentiles were chosen to reduce effects of vegetation activity reduction due to factors other than water stress (CWD). We then tested for each pixel whether the data support the model of a single linear decline of SIF (EF) with increasing CWD (Eq. 5), or, alternatively, a segmented regression model with one or two change points. The R package *segmented* was used [47]. The model with the lowest BIC was chosen and *S*_dSIF_ and *S*_dEF_ were quantified only for pixels where no significant change point was detected and where the regression of EF (SIF) vs. CWD had a significantly negative slope. “Flattening” EF (SIF) vs. CWD relationships were identified where a significant change point was detected and where the slope of the second regression segment was significantly less negative (*p* = 0.05 of *t* -test) compared to the slope of the first segment.

### Rooting zone water storage capacity and rooting depth model

Following ref. [23], the rooting zone water storage capacity *S*_0_ is modelled based on the assumption that plants size their rooting depth such that the corresponding *S*_0_ is just large enough to maintain function under the expected maximum cumulative water deficit (CWD) occurring with a return period of *T* years. Magnitudes of CWD extremes are estimated by fitting an extreme value distribution (Gumbel) to the annual maximum CWD values for each pixel separately, using the *extRemes* R package [24]. Magnitudes of extremes with a given return period *T* are derived and yield *S*_CWDX*T*_. *S*_CWDX*T*_ are translated into an effective depth *z*_CWDX*T*_ using estimates of the plant-available soil water holding capacity, based on soil texture data from a gridded version of the Harmonized World Soil Database [20, 72] and pedo-transfer functions derived by [4]. A global map of *z*_CWDX80_ is provided in Fig. S13.

### Estimating return periods

Diagnosed values of *S*_dSIF_ and *S*_dEF_ provide a constraint on the return period *T*. To yield stable estimates of *T* and avoid effects of the strong non-linearity of the function to derive *T* from the fitted extreme value distributions and magnitudes estimated by *S*_dSIF_ and *S*_dEF_, we pooled estimates *S*_dSIF_ (*S*_dEF_) and *S*_CWDX*T*_ values within 1° pixels (≤400 values). A range of discrete values *T* was screened (10, 20, 30, 40, 50, 60, 70, 80, 90, 100, 150, 200, 250, 300, 350, 400, 450, 500 years) and the best estimate *T* was chosen based on comparison to *S*_dSIF_ (*T*_SIF_) and to *S*_dEF_ (*T*_EF_), i.e., where the absolute value of the median of the logarithm of the bias was minimal.

### Rooting depth observations

The rooting depth data set (*N* = 5524) was compiled by combining and complementing published datasets from [54] and [18]. The data includes observations of the maximum rooting depth of plants taken from 361 published studies plus additional environmental and climate data. *z*_*r*_ was taken as the plant’s maximum rooting depth. Data were aggregated by sites (*N* = 359) based on longitude and latitude information. Sites were classified into biomes using maps of terrestrial ecoregions [49]. Quantiles (10%, 90%) were determined for each biome. For a subset of the data, where parallel measurements of the water table depth (WTD) was available, we conducted the same analysis, but took the minimum of WTD and *z*_*r*_.

## Results

### Estimating ET

Unbiased estimates of ET during rain-free periods are essential for determining CWD and estimating *S*_0_ and implied *z*_*r*_. We tested different remote sensing-based ET products and found that the ALEXI-TIR product, which is based on thermal infrared remote sensing [3, 27], exhibits no systematic bias during progressing droughts (Fig. S1). The stability in ET estimates from ALEXI-TIR during drought are enabled by its effective use of information about the surface energy partitioning, allowing inference of ET rates without reliance on a priori specified and inherently uncertain surface conductances [45] or shapes of empirical water stress functions [19], and without assumptions of rooting depth or effective *S*_0_. ALEXI-TIR is thus well-suited for estimating actual ET behaviour during drought without introducing circularity in inferring *S*_0_.

### Diagnosing *S*_0_ from SIF and EF

By employing first principles for the constraint of rooting zone water availability on ET and photosynthesis [66], we developed a method to derive how the sensitivity of these fluxes to increasing CWD relates to *S*_0_ and how this sensitivity can be used to reveal effects of access to extensive deep water stores (see Methods). Following this method, we determined the apparent *S*_0_ as the CWD at which vegetation “activity” ceases. The parallel information of evapotranspiration, precipitation, and snow mass balance enables a quantification of CWD over time. Vegetation activity was estimated from two alternative approaches: from the evaporative fraction (EF, defined as ET over net radiation), and from sun-induced fluorescence (SIF, normalised by incident shortwave radiation). SIF is a proxy for ecosystem photosynthesis [75] and is taken here from a spatially downscaled data product [16] based on GOME-2 data [33, 36]. Since net radiation and shortwave radiation are first-order controls on ET and SIF, respectively, and to avoid effects by seasonally varying radiation inputs, we used EF instead of ET, and considered the ecosystem-level fluorescence yield, quantified as SIF divided by shortwave radiation (henceforth referred to as ‘SIF') for all analyses.

Fig. 1 reveals large global variations in *S*_0_. Estimates based on EF and SIF agree closely (*R*^2^=0.78, Fig. S2). The lowest sensitivity of vegetation activity to an increasing CWD, and thus the largest apparent *S*_0_, is found in regions with a strong seasonality in radiation and water availability and substantial vegetation cover - particularly in monsoonal climates. In contrast, the lowest *S*_0_ values appear not only in regions where seasonal water deficits are limited due to short inter-storm duration (e.g., inner tropics) and/or low levels of potential evapotranspiration (e.g., high latitudes), but also in deserts and arid grasslands. This reflects the limited water surplus accumulating during rain events from which vegetation can draw during dry periods. A rapid decline of ET and SIF with an increasing CWD is related to vegetation cover dynamics, governed by greening after rain pulses and browning during dry periods [35].

**Figure 1.**
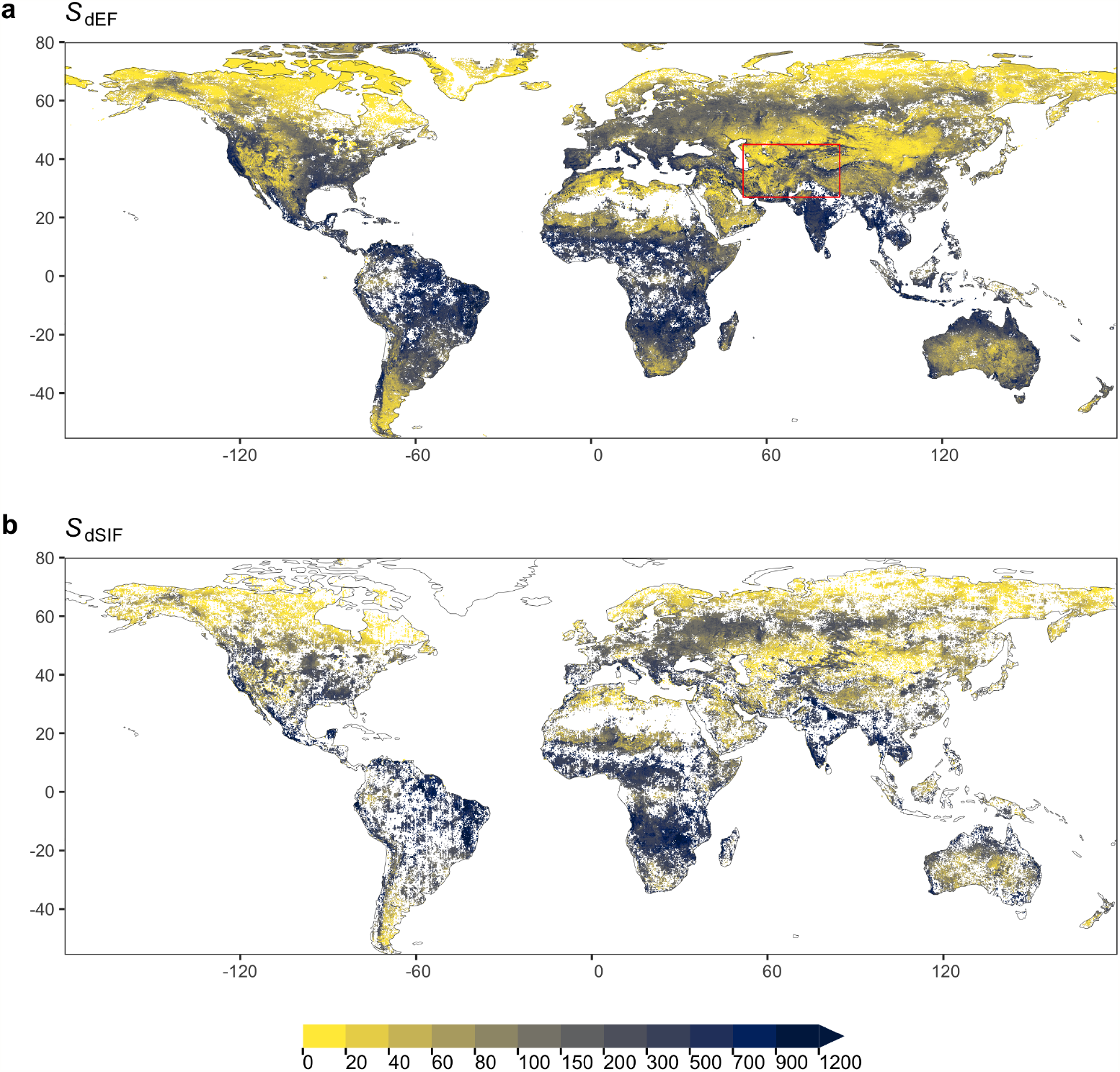
Rooting zone water storage capacity (mm) estimated from the sensitivity of the evaporative fraction (*S*_dEF_, a) and sun-induced fluorescence (*S*_dSIF_, b) to the cumulative water deficit (CWD). Data is aggregated to 0.1° resolution. The red box in (a) shows the outline of the magnified map provided in Fig. 2. Blank cells mark areas where all native cells at 0.05° resolution did not exhibit a significant and single, linearly declining relationship with increasing CWD.

Clear patterns emerge also at smaller scales (Fig. 2). *S*_dSIF_ and *S*_dEF_ consistently (Fig. S6) reveal how the sensitivity of photosynthesis and transpiration to drought stress varies across different topographical settings, indicating generally larger *S*_0_ in mountain regions and along rivers (Amu Darya) and deltas (Volga). We note however, that ALEXI ET estimates over mountainous terrain may be biased high where low incident net radiation and surface temperatures are caused not by high evaporative fractions but rather by topography effects and local shading. The maps of *S*_dSIF_ and *S*_dEF_ also bear strong imprints of human land use. Major irrigated cropland areas are congruent with some of the highest apparent *S*_0_ values recorded here. In these areas, our analysis yields particularly high CWD values and a low sensitivity of SIF and EF to CWD based on remotely sensed apparent ET and precipitation time series, without using information about the location and magnitude of irrigation. Other major irrigated areas appear as blank cells in Fig. 2 because the algorithm used to calculate CWD (see Methods) fails due to a long-term imbalance between *P* and ET and a “runaway CWD”. This indicates a sustained over-use of local water resources, caused by lateral water redistribution at scales beyond ~5 km, either via streamflow diversion or groundwater flow and extraction.

**Figure 2.**
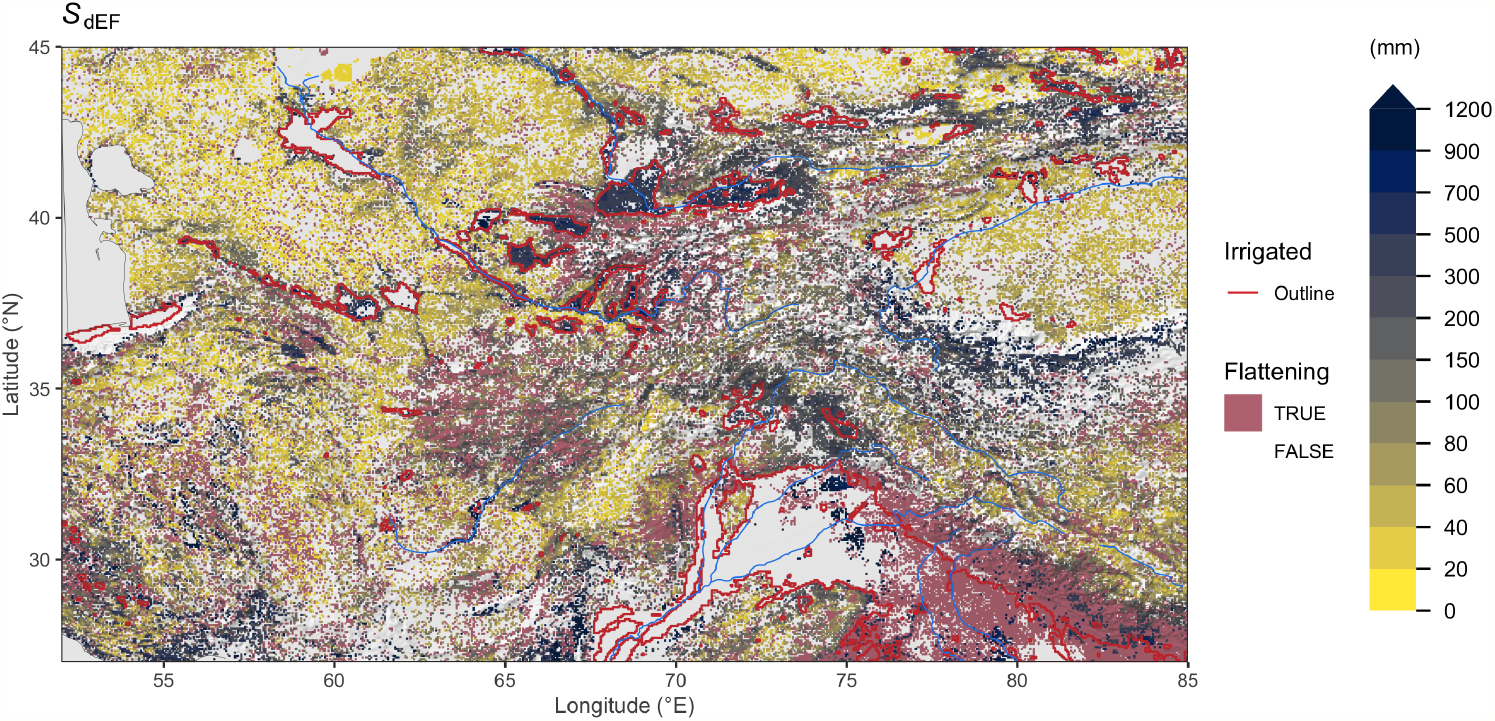
Rooting zone water storage capacity in Central Asia, estimated from the evaporative fraction (*S*_dEF_). Mauve areas (“flattening”) show grid cells where a significant reduction in the slope in EF vs. CWD was identified beyond a certain threshold. *S*_dEF_ values are not calculated for gridcells classified as “Flattening”. Red lines show outlines of major irrigated areas, i.e. where the irrigated land area fraction is above 30% [61]. Information about irrigated areas was used only for mapping here, but is not used for other parts of the analysis. Blank grid cells indicate areas with a sustained imbalance of ET being greater than *P*. Additional regional maps are provided by Figs. S3-S5.

Regressing vegetation activity against CWD also identifies locations where a decoupling of the two variables appears, i.e., where the sensitivity EF or SIF significantly decreases beyond a certain CWD threshold (“Flattening” in Fig. 2, see Methods). Such areas are particularly common in the vicinity of mountain regions, in areas with irrigated croplands, and in savannahs (Fig. S7). Related mechanisms may be at play. A flattening of the EF (SIF) vs. CWD relationship is likely due to the different portions of the vegetation having access to distinct water resources and respective storage capacities. In areas with large topographic gradients, this may be due to within-gridcell heterogeneity in plant access to the saturated zone. In savannahs, a shift in ET contributions from grasses and trees and a related shift in transpiration occurs as grasses, which are often more shallow-rooted than trees, senesce. In irrigated cropland areas, the flattening likely reflects land use heterogeneity within ~5 km grid cells and the persistent water access of irrigated fields while EF and SIF are reduced more rapidly in surrounding vegetation.

### Inferring *S*_0_ and *z*_*r*_ from CWD distributions

What controls spatial variations in *S*_0_ and *z*_*r*_ and the sensitivity of vegetation activity to water stress? Following ref. [23], we hypothesized that plants are adapted to the local hydroclimate and size their *z*_*r*_ to maintain transpiration and photosynthesis in the face of water deficits during dry periods commonly experienced over the course of a plant’s lifetime. Specifically, we hypothesized that *S*_0_ can be inferred from the distribution of annual CWD maxima at a given location and that *S*_0_ corresponds to a CWD extreme recurring with a return period of *T* years, where *T* is on the same order of magnitude as the typical lifetime of dominant plants (see Methods). We start by using *T* = 80 years for estimating *S*_0_ and *z*_*r*_ here (termed *S*_CWDX80_ and *z*_CWDX80_) and fit CWD extreme value distributions separately for each pixel.

Fig. 3a shows the global distribution of *S*_CWDX80_ and reveals patterns across multiple scales - in close agreement with *S*_dSIF_ and *S*_dEF_ (*R*^2^ = 0.76 and *R*^2^ = 0.84, respectively, Fig. S6). This indicates that the sensitivity of vegetation activity to an increasing CWD is strongly controlled by the magnitude of local CWD extremes. Perennially moist tropical regions around the Equator (northwestern Amazon, Congo, southeast Asia) where persistent cloud cover prohibits the derivation of *S*_dSIF_ and *S*_dEF_, emerge here as characterised by particularly low *S*_CWDX80_. Fine granularity and large spatial heterogeneity of *S*_CWDX80_ at regional scales, particularly in arid and semi-arid regions, reveal the importance of the local topographical setting (and irrigation, see Fig. 2) for determining plant-available water storage capacities (Figs. S8-S11). Variations are likely to extend to even smaller scales along the hillslope topography [18] and within individual forest stands [1]. These scales lie beyond the resolution of the satellite remote sensing data used here to calculate CWD.

**Figure 3.**
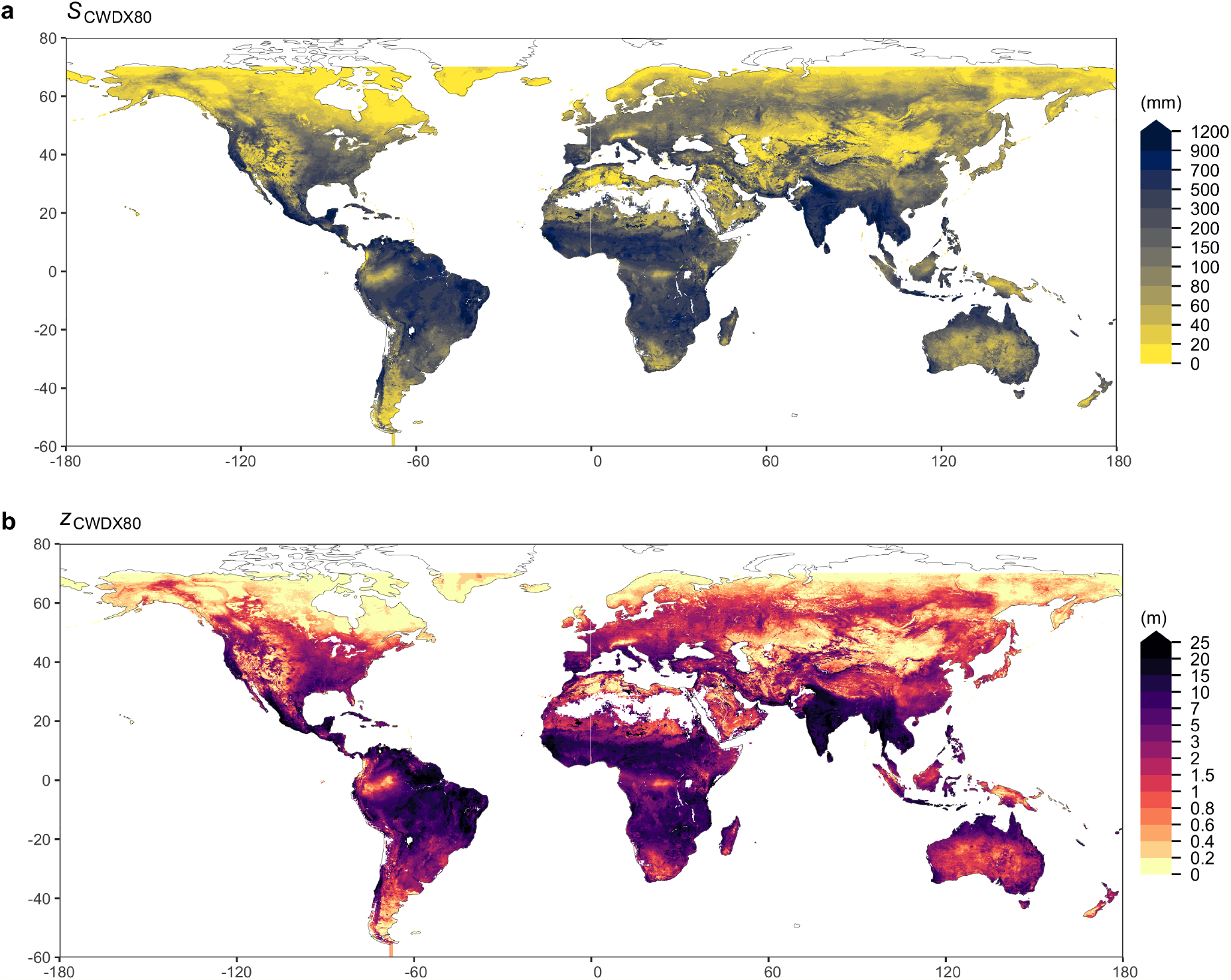
Return periods *T* (yr), diagnosed from EF (a) and SIF (b). To diagnose *T*, a range of alternative values of *T* are screened and the corresponding range of values *S*_CWDX*T*_ are compared to *S*_dEF_ (*S*_dSIF_) within 1° grid cells (resolution of maps shown here). The best matching *T* was retained for each gridcell, yielding a global distribution of *T*_EF_ (*T*_SIF_). The lower panel shows the distribution of diagnosed return periods *T*

### Validation with rooting depth observations

Next, we investigate whether the strong control of CWD extremes on the sensitivity of vegetation activity to water deficits reflects the adaptation of plant rooting depth to the hydroclimate. For this, we compare *S*_CWDX80_ against (fully independent) field observations of *z*_*r*_ from a novel compilation of tree-level data with global coverage, extended from previous work [18, 56], now encompassing 5524 observations (see also Methods). Data was aggregated by taking the mean of individual observations for 1705 globally distributed sites (SI Fig. S12). To avoid the inevitable scale mis-match between in situ ecosystem observations and global remote sensing [39], we focused our evaluation against observed rooting depth at the scale of biomes. We converted estimates of *S*_CWDX80_ into a corresponding depth (*z*_CWDX80_) by accounting for soil texture along the rooting profile (see Methods, Fig. 3b).

**Figure 3.**
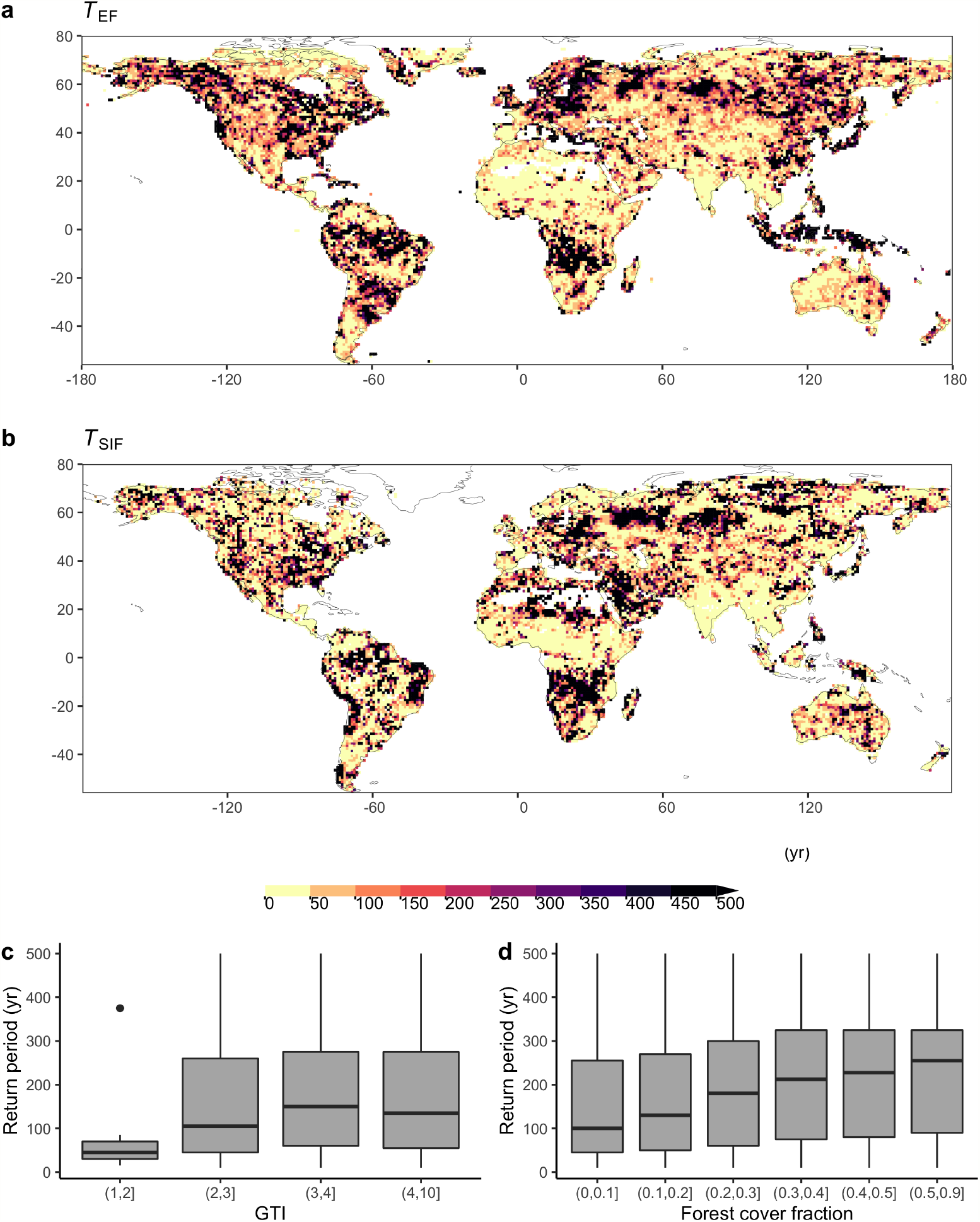
Spatial variations of (a) the rooting zone water storage capacity, estimated by *S*_CWDX80_ (mm), and (b) the apparent plant rooting depth *z*_CWDX80_ (m). Values are remapped to a 0.1° resolution. Blank grid cells are either permanent inland water bodies and ocean, or locations with long-term accumulation of water deficits. Values are removed in gridcells where more than 80% is non-vegetation surface based on MODIS Landcover [22].

Our rooting depth estimates *z*_CWDX80_ capture general differences in rooting depth across biomes (Fig. 4). Predicted and observed biome-level maximum rooting depth (90% quantiles) are in good agreement (Fig. 4c), while the lower (10%) quantiles appear to be overestimated by *z*_CWDX80_ (Fig. 4b). Using a subset of the data where information about the water table depth (WTD) is provided (490 entries from 359 sites), we limited predicted *z*_CWDX80_ to the value of the observed local WTD (53% of all observations). This yields an improved agreement (Fig. 4d) and suggests that our predictive model for *z*_*r*_ overestimates values where the roots access groundwater. While acting as a constraint on the rooting depth [18], plant access to groundwater implies sustained transpiration during dry periods, correspondingly large cumulative water deficits and, by implication of the model design, large *S*_CWDX80_ and *z*_CWDX80_. In other words, large inferred *S*_CWDX80_ and *z*_CWDX80_ reflect sustained transpiration, enabled not by the exceptional depth of roots, but by roots tapping into the groundwater, and CWD is limited not by the supply of water, but by the demand (effectively radiation-limited ET). This highlights that estimates of *S*_0_ and *z*_*r*_ presented here are to be interpreted as apparent quantities, subject to groundwater influence (and irrigation, as shown above). It also demonstrates that knowledge about the groundwater depth and whether vegetation taps into it is key for explaining ET during dry periods at a large proportion of sites investigated here.

**Figure 4.**
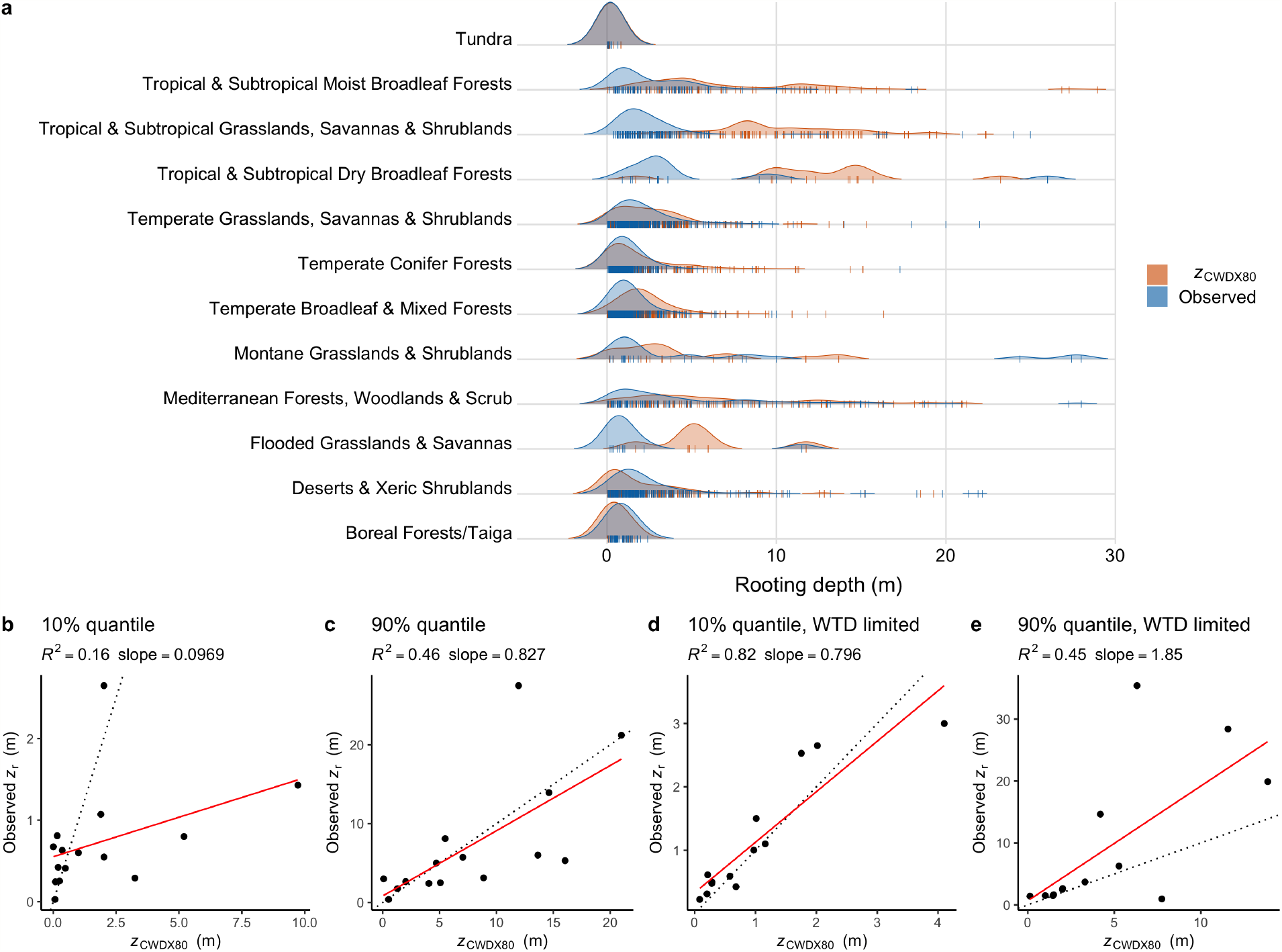
Modelled and observed rooting depth by biomes. (a) Kernel density estimates of observed and predicted (*z*_CWDX80_) rooting depth by biomes, based on data aggregated by sites, shown by vertical colored tick marks. 10% and 90% quantiles of observed vs. predicted (*z*_CWDX80_) rooting depth by biome of all data (b,c), and of a subset of the data where water table depth was measured along with rooting depth (d,e). Classification of sites into biomes was done based on [49]. Dotted lines in b-e represent the 1:1 line.

### Estimating return periods

Underlying the estimates of *S*_CWDX80_ is the assumption that plant rooting strategies are adapted to sustain CWD extremes with a return period *T* = 80 years. *S*_dSIF_ and *S*_dEF_ provide an independent constraint to test this assumption. Fig. 3 suggests that *T* is not a global constant and that a general pattern emerges, largely in consistency between the independent estimates *T*_EF_ and *T*_SIF_. A clear tendency towards higher *T* emerges with an increasing gridcell average forest cover fraction (Fig. 3d). *S*_0_ appears to be adapted to 500-year and even rarer events in forested regions but to lower *T* outside. A return period on the order of multiple centuries is consistent with the typical lifetime of trees [6] and suggests that optimal plant adaptation of life history strategies to a stochastic environment may be understood as being governed by frequencies of climate extremes in relation to the lifetime of affected organisms. Interestingly, substantial variations in *T* remain even within land cover types (e.g. the boreal forests of Russia). This variation may be related to the large-scale topographical setting and the tendency towards shallow groundwater table depths, as measured by the Compound Topographic Index ( [41], Fig. 3c). In such areas, vegetation appears to sustain particularly rare CWD extremes (diagnosed here from its low sensitivity to CWD during the observation period), possibly enabled by roots’ access to a relatively shallow saturated zone.

## Discussion and conclusion

*S*_0_ is a central quantity in determining land-atmosphere exchange and vegetation resistance to drought, but its spatial variations cannot be observed directly and variations within vegetation types are largely ignored in models. Using first-principles modelling and integrating multiple data streams at daily and ~5 km resolution, we diagnosed a hydrologically effective ecosystem-level *S*_0_ from the sensitivity of vegetation activity to CWD. Our analysis revealed extensive spatial heterogeneity of plant-available moisture storage in the Critical Zone, with apparent variations extending from the continental to the regional scale. While large-scale variations in *S*_0_ are mainly driven by the hydroclimate, more fine-grained variations within regions are linked to topography and how vegetation and land use is distributed across the landscape and determines actual ET.

By detecting where the sensitivity of vegetation activity to an increasing CWD declines beyond a certain CWD threshold, we identified locations where a portion of the (mean of *T*_EF_ and *T*_SIF_) within bins of the Compound Topographical Index [41] (c) and forest cover fractions (MOD44B [29]) (d). vegetation may have access to deep water stores and where groundwater may contribute to land-atmosphere coupling [42]. Traditional land surface models, which typically employ the assumption of free drainage out of the bottom soil layer and simulate continuously increasing water stress effects as CWD increases, may not be suitable for reliably simulating land-atmosphere exchange during prolonged dry periods in these regions.

Diagnosed values of *S*_0_ are representative of water stored across the entire rooting zone - from the soil surface to the weathered bedrock and groundwater. Particularly in regions with pronounced dry seasons, magnitudes of *S*_CWDX80_ greatly exceed typical values of the total soil water holding capacity when considering the top 1 or 2 meters of the soil column and texture information from global databases ( [20], Fig. S13) - as sometimes done in models [11, 28, 44]. Recent land surface model developments have included deeper soil layers, but effective *S*_0_ are subject also to choices of *z*_*r*_. The discrepancy in magnitude and spatial patterns of total 1 (2) m soil water holding capacity and *S*_0_ diagnosed here underlines the critical role of plant access to deep water and the need to extend the focus beyond moisture in the top 1-2 m of soil for simulating land-atmosphere exchange [12]. We note that using *S*_CWDX80_ (*z*_CWDX80_) for directly parameterizing *S*_0_ (*z*_*r*_) in global models may be misleading in forested areas with particularly small *S*_CWDX80_. Additional effects of how *z*_*r*_ determines access to belowground resources and function (e.g., nutrients, mechanical stability) should be considered.

Our estimate of *S*_0_ implicitly includes water intercepted by leaf and branch surfaces, internal plant water storage, and moisture stored in the topsoil and supplied to evaporation. These components are generally smaller in magnitude compared to moisture storage supplied to transpiration [64], and their contribution to ET declines rapidly as CWD increases. (Moreover, readily available global evaporation data is lacking [39]). Hence, spatial variations in *S*_0_ primarily reflect variations mediated by moisture stored across the root zone.

Our analysis identified mountain regions as “water towers” with particularly high *S*_0_, in spite of shallow soil and regolith depths along upland hillslopes [51]. This could be due to hillslope-scale variations in groundwater depth where lateral flow pushes the saturated zone closer to the surface in valley bottoms and enables sustained transpiration during prolonged rain-free periods. Lateral subsurface flow is not explicitly accounted for, but can be considered relatively small at the scale investigated here (~5 km). Nevertheless, local convergence (divergence) acts to supply (remove) moisture and sustain (reduce) ET, leading to larger (smaller) CWD values. This effect should be captured by the relatively accurate remote sensing of ET by TIR (Fig. S1), and likely contributes to the strong contrast in *S*_CWDX80_ along topographic gradients (Figs. S8-S11). However, further research should assess the accuracy of spatial variations in annual mean ET and potential effects of terrain, where lower temperature rise signals on shaded slopes may be mis-interpreted by the ALEXI algorithm as signatures of higher ET. Since ET data are used both in *S*_CWDX80_ and in *S*_dEF_ and *S*_dSIF_, these quantities are not entirely independent, potentially leading to an overestimation of their correlation strengths (Fig. S7). Nevertheless, our predictions of continental-scale variations in *S*_0_ and implied *z*_*r*_ agree well with patterns derived from an extended dataset of tree-level *z*_*r*_ observations - an entirely independent observational constraint.

In contrast to earlier studies [18, 34, 39, 54, 74], here we mainly focused on *S*_0_, instead of *z*_*r*_. This makes our quantification independent of large variations in porosity along depths extending beneath the soil and focuses on the quantity relevant for the surface water balance. Nevertheless, where vegetation accesses the saturated zone, effective *S*_0_ may be even larger than numbers derived here based on past CWD time series. Similarly, the concept of a constrained water storage capacity is inappropriate where irrigation supplies water to ET. For comparison to observations and to earlier studies, we converted *S*_0_ into *z*_*r*_ using soil texture information provided for the top 1 m of the soil profile [20, 72]. This was chosen due to a lack of information about porosity of the subsoil.

Our *z*_*r*_ map (Fig. S13) yields patterns that are mostly consistent with previous estimates of the global distribution of *z*_*r*_ [18, 34, 39, 54, 74], but differ in important aspects. Deepest roots are predicted here in monsoonal climates with substantial vegetation cover, particularly in south and southeast Asia, but are predicted along the arid edges of vegetated zones elsewhere [18, 74]. This may be due to differences between TIR-derived ET versus ET derived from other approaches used in [18, 74]. Our evaluation against observations suggests that the distribution of *z*_*r*_ is reliably predicted by *z*_CWDX80_ for deserts and xeric shrublands and in tropical and subtropical broadleaf forests (Fig. 4). Suggested deep rooting in dry steppe regions of Kazakhstan [18, 74] and further east [39, 54] are not corroborated here. Mismatches may be due to the rapid dynamics of green foliage cover and ensuing water losses in these regions. In contrast to other studies [74], the ET and *P* time series used here capture these dynamics at a daily resolution. Common to most [18, 34, 54, 69, 74], but not all [39], global *z*_*r*_ estimates, including the present one, is the characteristic contrast between *z*_*r*_ in the northwestern and northeastern Amazon. Our estimates of *S*_CWDX80_ are largely consistent with results by [69] who employed a very similar approach and provided estimates at 0.5° resolution. However, highest values estimated here are substantially higher and distributed over a larger area, particularly in South Asia.

Taken together, constraints available from local *z*_*r*_ observations and from global remote sensing of vegetation activity converge on consistent patterns across multiple spatial scales and suggest that belowground vegetation structure can be sensed from space. Using methods developed here, combined with emerging data from recently launched satellite missions for thermal infrared remote sensing (e.g., ECOSTRESS) [73] bears promise for resolving *S*_0_ variations at the hillslope scale (100 m - 1 km). This will provide critical information for reliably estimating water deficits and spatial heterogeneity of ET across the landscape at scales relevant to multiple stakeholders. Contrasting future CWD extremes with their past distribution will yield insight into the severity of droughts with respect to levels to which *z*_*r*_ is adapted today.

Our study revealed a tight control of the climatology of water deficits on vegetation sensitivity to drought stress, and demonstrated how land-atmosphere interactions and the Critical Zone water storage capacity are governed by vegetation and its adaptation to the hydroclimate and topography across the globe. It remains to be shown whether plasticity in *z*_*r*_ is sufficiently rapid to keep pace with a changing climate with strong and wide-spread increases in rainfall variability [52], and to what degree rising CO_2_ alters plant water use and their carbon economy and thereby the costs and benefits of deep roots [15, 31].

## Acknowledgments

B.D.S was funded by the Swiss National Science Foundation grant no. PCEFP2 181115 and by the generosity of Eric and Wendy Schmidt by recommendation of the Schmidt Futures program. We acknowledge ICDC, CEN, University of Hamburg for data support. SJTD was funded by the NSF GRFP and LTER, and the NASEM Ford Foundation Fellowship Program. AGK was supported by NSF DEB-1942133.

