## Supplementary Information for "Global distribution of the rooting zone water storage capacity reflects plant adaptation to the environment"

#### S1 Evaluating ET datasets

Unbiased estimates of ET during rain-free periods are essential for implying rooting depth following the method employed here. Various global ET data products are available. The widely-used Priestly-Taylor (PT) model uses net radiation for estimating the energy supply available for evapotranspiration. Additional empirical scalars have to be applied to reduce ET estimates under water-limited conditions. The Penman-Monteith (PM) model is conceptually related to the PT model, but explicitly resolves effects of atmospheric dryness (vapour pressure deficit) and multiple conductance terms that control ET. As for the PT model, water stress effects have to be factored in additionally, commonly by reducing the surface conductance to transpiration.

The challenge with both PT and PM models is that water stress factors themselves rely, either directly or indirectly, on assumptions regarding plant rooting depth and are thus not suitable for use with the methods applied here. They would introduce circular reasoning and the implied rooting depth would directly reflect the assumptions regarding rooting depth or sensitivity of conductance to water stress, introduced in the PT and PM models themselves. Moreover, if rooting depth assumptions (and thus assumptions of effective S0) are inaccurate, respective ET estimates should exhibit a systematic bias related to the severity of water stress. A potential solution to this problem is to rely on ET products that make no a priori assumptions regarding rooting depth. Thermal infrared (TIR) -based methods rely primarily on land surface and air properties for estimating ET. However, previous studies found generally larger scatter in TIR-based ET estimates. For the present analysis, it's particularly important that estimates exhibit no systematic bias and that the bias is not related to the duration of rain-free periods and the severity of water stress. Therefore, we first tested a set of ET-based datasets with global coverage, before using the data for the present analysis.

We evaluated the bias of modelled versus observed ET, measured with the eddy covariance technique. The data were provided through the FLUXNET 2015 Tier 1 dataset Pastorello et al. (2021). To focus the evaluation on model performance under water stress, we subset the data to sites and periods where clear effects of water stress on photosynthetic light use efficiency have been identified Stocker et al. (2018). Data were aligned by the onset of periods with apparent water stress effects (droughts) to determine the "day into drought" (dday). Data were then normalised to pre-drought levels, separately for each site, and aggregated across drought events and years. Finally, we calculated quantiles of the normalised, aggregated bias versus

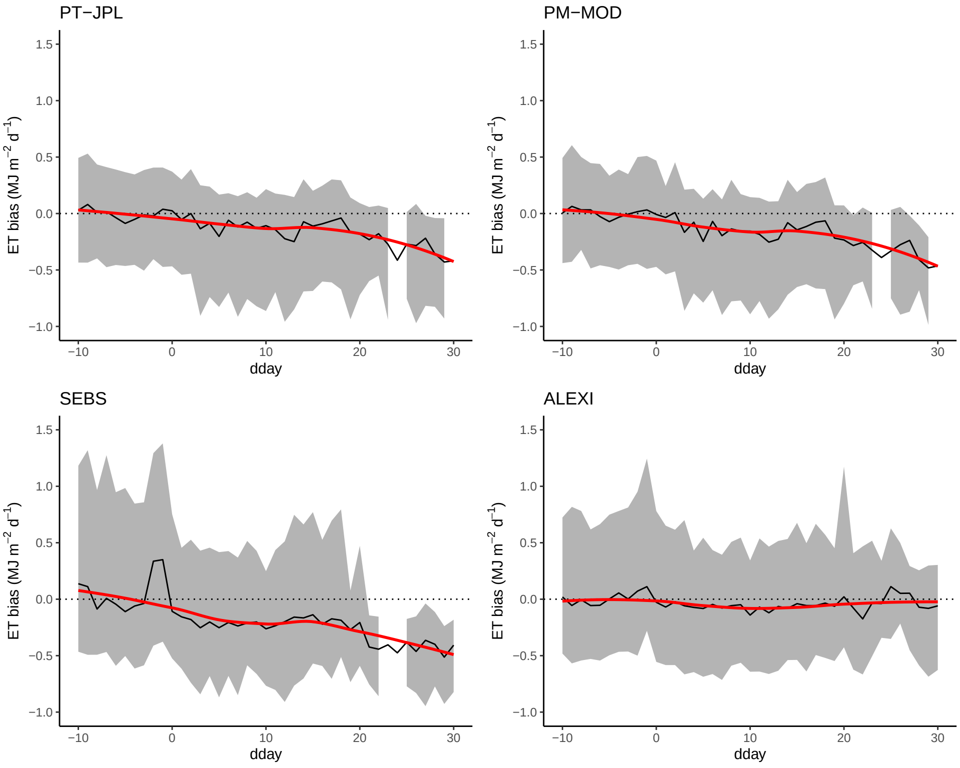


**Figure S1:** Bias of modelled versus observed ET based on four different algorithms applied to remote sensing data, and evaluated with observations from ecosystem flux measurements (FLUXNET Eddy covariance) collected during periods of drought. The black lines indicate the median bias for each day into the drought ('dday'), derived from multiple drought events and sites. The shaded area expands from the 33 to the 66 percentile, the red line is a LOESS smoothing line based on the median.

dday.

Fig. S1 reveals that indeed, although exhibiting less scatter before, the PT and PM-based algorithms perform less reliably than the TIR-based algorithm during rain-free periods. ALEXI-TIR provides accurate estimates of ET with no systematic bias related to the duration of droughts and the severity of water stress, and provides a robust observation of surface water loss. Since ALEXI-TIR makes no assumptions regarding rooting depth or effective S0, and in view of its robust performance under water-stressed conditions, we apply this data product for all analyses shown here.

### S2 Supplementary figures

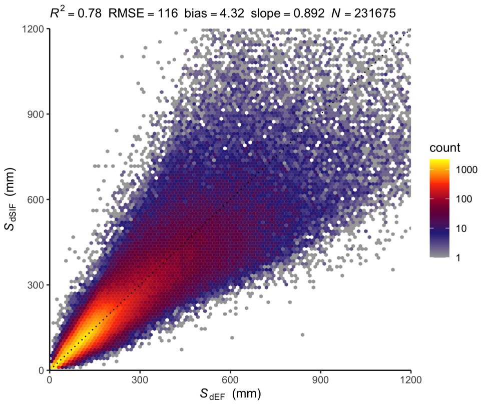


**Figure S2:** Correlation of $S_0$, diagnosed from EF ($S_{\mathrm{dEF}}$) and from SIF ($S_{\mathrm{dSIF}}$). Metrics of the correlation are given by text annotation on top of the figures (RMSE is the root mean square error). The dashed black line indicates the 1:1 line.

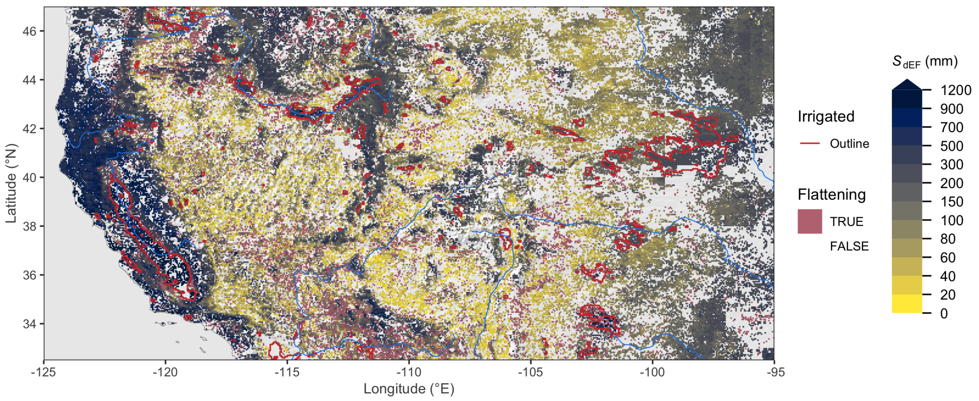


**Figure S3:** Rooting zone water storage capacity in the Western USA, estimated from the evaporative fraction ($S_{dEF}$). Mauve areas ("flattening") show grid cells where a significant reduction in the slope in EF vs. CWD was identified beyond a certain threshold. Red lines show outlines of major irrigated areas, i.e. where the irrigated land area fraction is above 30% Siebert et al. (2005). Blank grid cells indicate areas with a sustained imbalance of ET being greater than P.

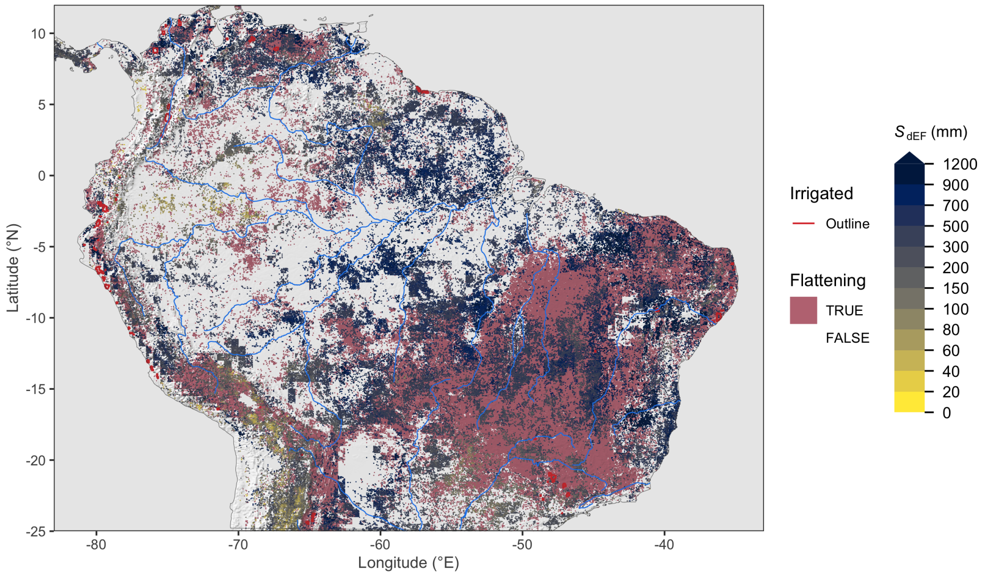


**Figure S4:** Rooting zone water storage capacity in the Amazon region, estimated from the evaporative fraction ($S_{dEF}$). Mauve areas ("flattening") show grid cells where a significant reduction in the slope in EF vs. CWD was identified beyond a certain threshold. Red lines show outlines of major irrigated areas, i.e. where the irrigated land area fraction is above 30% Siebert et al. (2005). Blank grid cells indicate areas with a sustained imbalance of ET being greater than P.

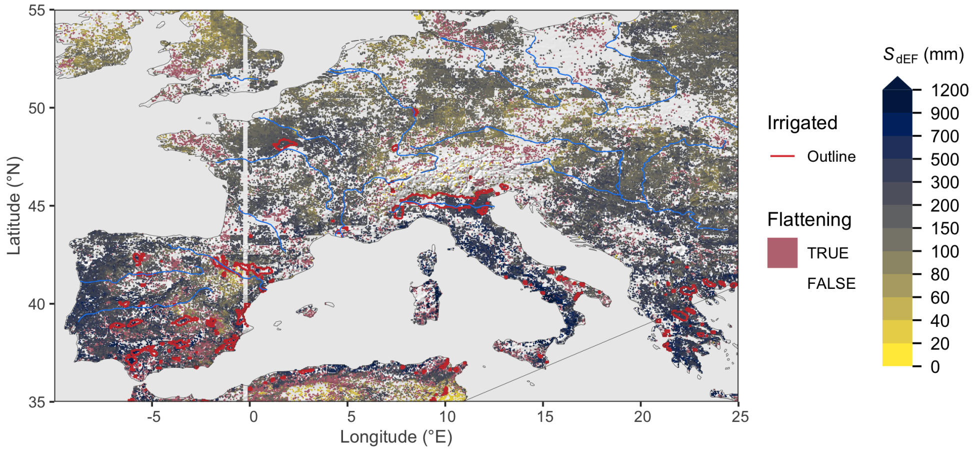


**Figure S5:** Rooting zone water storage capacity in Europe, estimated from the evaporative fraction ($S_{dEF}$). Mauve areas ("flattening") show grid cells where a significant reduction in the slope in EF vs. CWD was identified beyond a certain threshold. Red lines show outlines of major irrigated areas, i.e. where the irrigated land area fraction is above 30% Siebert et al. (2005). Blank grid cells indicate areas with a sustained imbalance of ET being greater than P.

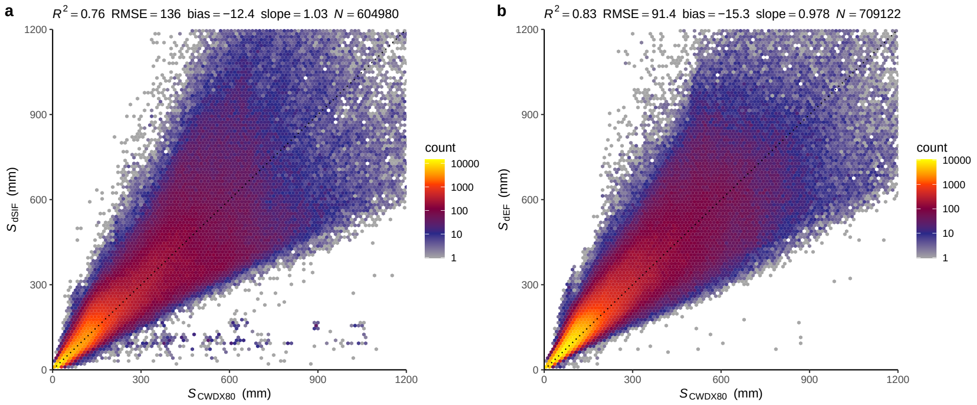


**Figure S6:** Correlation of $S_{\text{dEF}}$ (a) and $S_{\text{dSIF}}$ (b) with $S_{\text{CWDX80}}$. Metrics of the correlation are given by text annotation on top of the figures (RMSE is the root mean square error). The dashed black line indicates the 1:1 line.

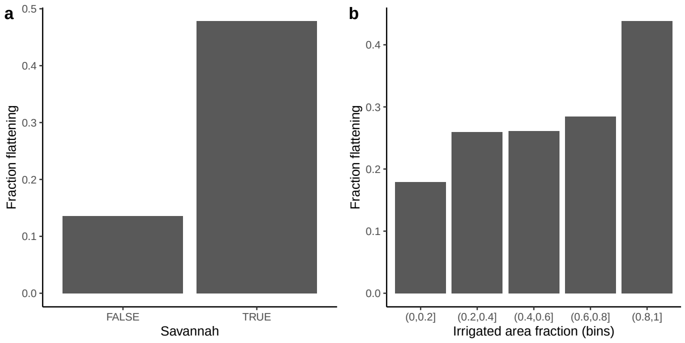


**Figure S7:** Fraction of grid cells where a flattening relationship between EF and CWD was detected (see Methods), in savannahs and outside (a) and depending on the irrigated area fraction (b). 'Savannah' is taken as 'Woody Savannah' from MODIS MCD12C1 for year 2010 Friedl and Sulla-Menashe (2015). The fraction of irrigated areas is from Siebert et al. (2005), taken as actually irrigated areas as a fraction of land area.

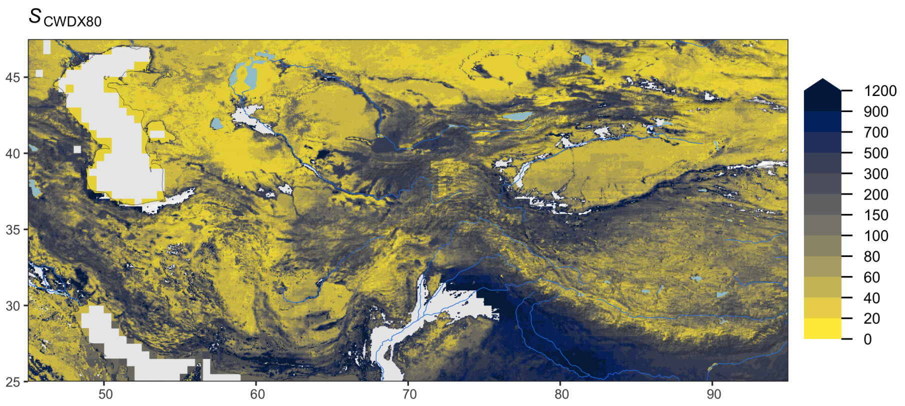


**Figure S8:** Rooting zone water storage capacity in Central Asia, estimated as cumulative water deficit extreme events with a return period of 80 years ($S_{CWDX80}$). Blank grid cells indicate areas with a sustained imbalance of ET being greater than $P$. Grey shading shows the surface topography, light blue lines and areas represent major rivers and lakes.

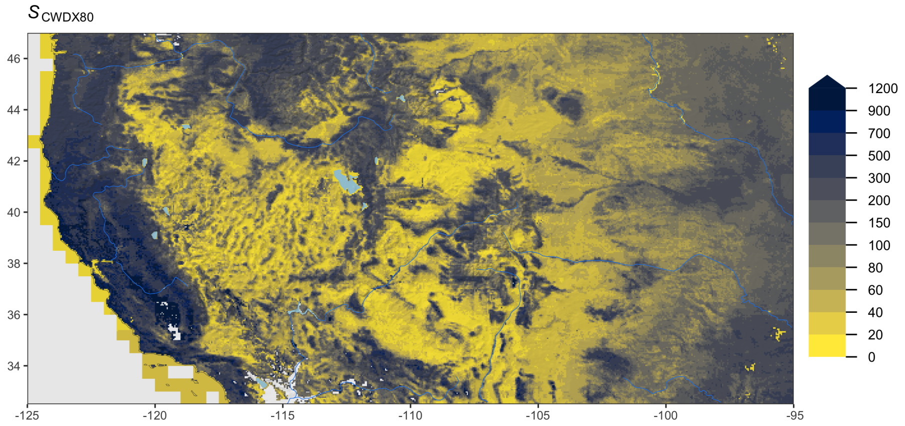


**Figure S9:** Rooting zone water storage capacity in the western USA, estimated as cumulative water deficit extreme events with a return period of 80 years ($S_{\mathrm{CWDX80}}$). Blank grid cells indicate areas with a sustained imbalance of ET being greater than $P$. Grey shading shows the surface topography, light blue lines and areas represent major rivers and lakes.

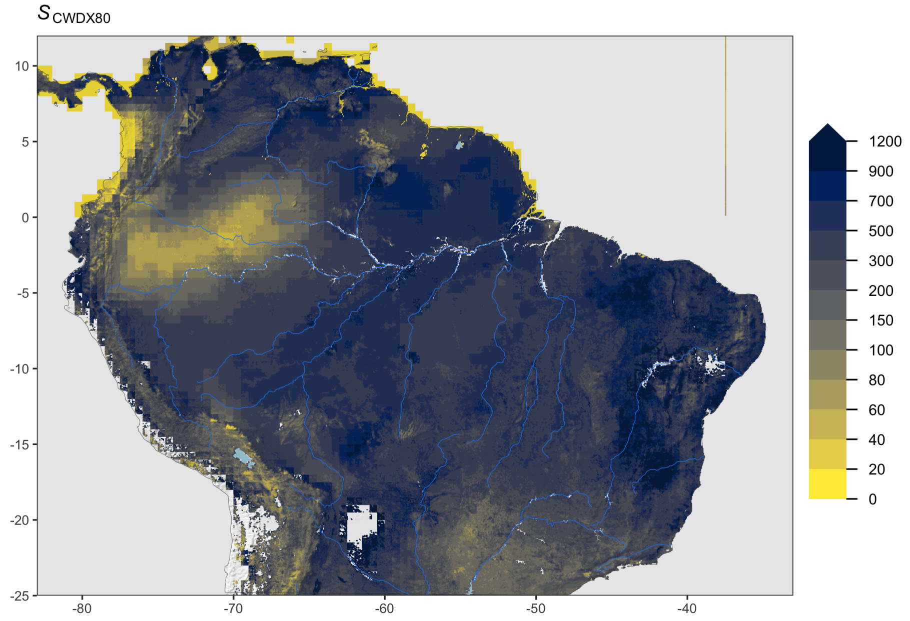


**Figure S10:** Rooting zone water storage capacity in the Amazon region, estimated as cumulative water deficit extreme events with a return period of 80 years ($S_{\text{CWDX80}}$). Blank grid cells indicate areas with a sustained imbalance of ET being greater than $P$. Grey shading shows the surface topography, light blue lines and areas represent major rivers and lakes.

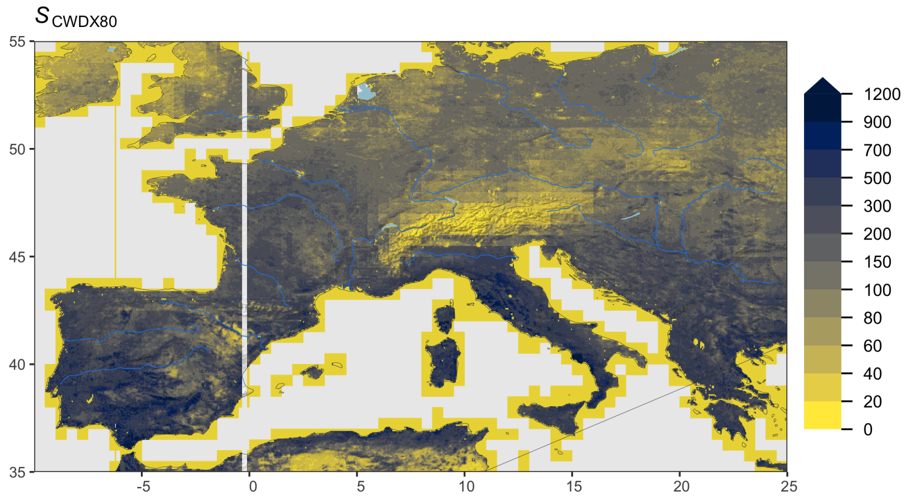


**Figure S11:** Rooting zone water storage capacity in Europe, estimated as cumulative water deficit extreme events with a return period of 80 years ($S_{\mathrm{CWDX80}}$). Blank grid cells indicate areas with a sustained imbalance of ET being greater than $P$. Grey shading shows the surface topography, light blue lines and areas represent major rivers and lakes.

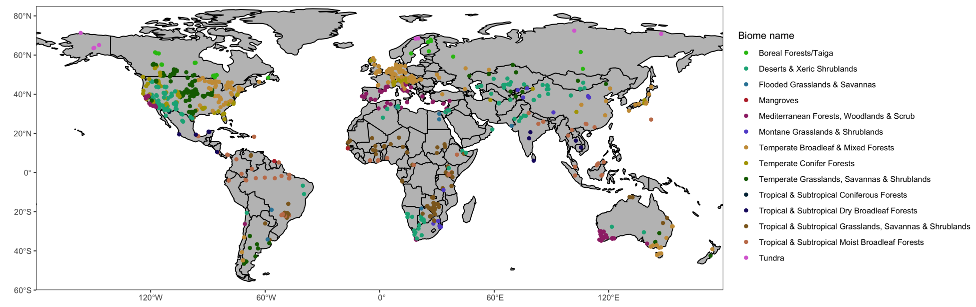


**Figure S12:** Location and biome type of sites where rooting depth data is provided and used here for comparison against $z_{\text{CWDX80}}$. Biome classification was done based on Olson et al. (2001).

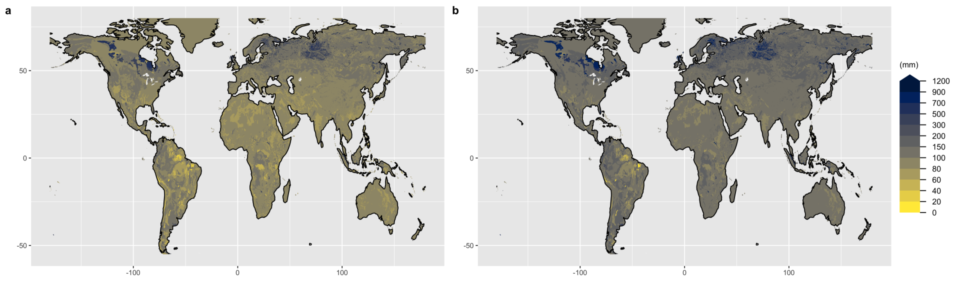


**Figure S13:** Integrated soil water holding capacity across the top 1 m (a) and the top 2 m (b). Values are calculated based on soil texture information from the HWSD **?** and pedo-transfer functions based on Balland et al. (2008). HWSD provides information for a top layer (0-30 cm depth) and a bottom layer (30-100 cm depth). For the top 2 m shown in (b), we assumed values from the bottom layer for 100-200 cm depth.

Pastorello, G., Trotta, C., Canfora, E., Chu, H., Christianson, D., Cheah, Y.-W., Poindexter, C., Chen, J., Elbashandy, A., Humphrey, M., Isaac, P., Polidori, D., Reichstein, M., Ribeca, A., van Ingen, C., Vuichard, N., Zhang, L., Amiro, B., Ammann, C., Arain, M. A., Ardö, J., Arkebauer, T., Arndt, S. K., Arriga, N., Aubinet, M., Aurela, M., Baldocchi, D., Barr, A., Beamesderfer, E., Marchesini, L. B., Bergeron, O., Beringer, J., Bernhofer, C., Berveiller, D., Billesbach, D., Black, T. A., Blanken, P. D., Bohrer, G., Boike, J., Bolstad, P. V., Bonal, D., Bonnefond, J.-M., Bowling, D. R., Bracho, R., Brodeur, J., Brümmer, C., Buchmann, N., Burban, B., Burns, S. P., Buysse, P., Cale, P., Cavagna, M., Cellier, P., Chen, S., Chini, I., Christensen, T. R., Cleverly, J., Collalti, A., Consalvo, C., Cook, B. D., Cook, D., Coursolle, C., Cremonese, E., Curtis, P. S., D'Andrea, E., da Rocha, H., Dai, X., Davis, K. J., De Cinti, B., de Grandcourt, A., De Ligne, A., De Oliveira, R. C., Delpierre, N., Desai, A. R., Di Bella, C. M., di Tommasi, P., Dolman, H., Domingo, F., Dong, G., Dore, S., Duce, P., Dufrêne, E., Dunn, A., Dušek, J., Eamus, D., Eichelmann, U., ElKhidir, H. A. M., Eugster, W., Ewenz, C. M., Ewers, B., Famulari, D., Fares, S., Feigenwinter, I., Feitz, A., Fensholt, R., Filippa, G., Fischer, M., Frank, J., Galvagno, M., Gharun, M., Gianelle, D., Gielen, B., Gioli, B., Gitelson, A., Goded, I., Goeckede, M., Goldstein, A. H., Gough, C. M., Goulden, M. L., Graf, A., Griebel, A., Gruening, C., Grünwald, T., Hammerle, A., Han, S., Han, X., Hansen, B. U., Hanson, C., Hatakka, J., He, Y., Hehn, M., Heinesch, B., Hinko-Najera, N., Hörtnagl, L., Hutley, L., Ibrom, A., Ikawa, H., Jackowicz-Korczynski, M., Janouš, D., Jans, W., Jassal, R., Jiang, S., Kato, T., Khomik, M., Klatt, J., Knohl, A., Knox, S., Kobayashi, H., Koerber, G., Kolle, O., Kosugi, Y., Kotani, A., Kowalski, A., Kruijt, B., Kurbatova, J., Kutsch, W. L., Kwon, H., Launiainen, S., Laurila, T., Law, B., Leuning, R., Li, Y., Liddell, M., Limousin, J.-M., Lion, M., Liska, A. J., Lohila, A., López-Ballesteros, A., López-Blanco, E., Loubet, B., Loustau, D., Lucas-Moffat, A., Lüers, J., Ma, S., Macfarlane, C., Magliulo, V., Maier, R., Mammarella, I., Manca, G., Marcolla, B., Margolis, H. A., Marras, S., Massman, W., Mastepanov, M., Matamala, R., Matthes, J. H., Mazzenga, F., McCaughey, H., McHugh, I., McMillan, A. M. S., Merbold, L., Meyer, W., Meyers, T., Miller, S. D., Minerbi, S., Moderow, U., Monson, R. K., Montagnani, L., Moore, C. E., Moors, E., Moreaux, V., Moureaux, C., Munger, J. W., Nakai, T., Neirynck, J., Nesic, Z., Nicolini, G., Noormets, A., Northwood, M., Nosetto, M., Nouvellon, Y., Novick, K., Oechel, W., Olesen, J. E., Ourcival, J.-M., Papuga, S. A., Parmentier, F.-J., Paul-Limoges, E., Pavelka, M., Peichl, M., Pendall, E., Phillips, R. P., Pilegaard, K., Pirk, N., Posse, G., Powell, T., Prasse, H., Prober, S. M.,

Rambal, S., Rannik, Ü., Raz-Yaseef, N., Rebmann, C., Reed, D., de Dios, V. R., Restrepo-Coupe, N., Reverter, B. R., Roland, M., Sabbatini, S., Sachs, T., Saleska, S. R., Sánchez-Cañete, E. P., Sanchez-Mejia, Z. M., Schmid, H. P., Schmidt, M., Schneider, K., Schrader, F., Schroder, I., Scott, R. L., Sedlák, P., Serrano-Ortíz, P., Shao, C., Shi, P., Shironya, I., Siebicke, L., Šigut, L., Silberstein, R., Sirca, C., Spano, D., Steinbrecher, R., Stevens, R. M., Sturtevant, C., Suyker, A., Tagesson, T., Takanashi, S., Tang, Y., Tapper, N., Thom, J., Tomassucci, M., Tuovinen, J.-P., Urbanski, S., Valentini, R., van der Molen, M., van Gorsel, E., van Huissteden, K., Varlagin, A., Verfaillie, J., Vesala, T., Vincke, C., Vitale, D., Vygodskaya, N., Walker, J. P., Walter-Shea, E., Wang, H., Weber, R., Westermann, S., Wille, C., Wofsy, S., Wohlfahrt, G., Wolf, S., Woodgate, W., Li, Y., Zampedri, R., Zhang, J., Zhou, G., Zona, D., Agarwal, D., Biraud, S., Torn, M., and Papale, D.: Author Correction: The FLUXNET2015 dataset and the ONEFlux processing pipeline for eddy covariance data, Sci Data, 8, 72, 2021.

Siebert, S., Döll, P., Hoogeveen, J., Faures, J.-M., Frenken, K., and Feick, S.: Development and validation of the global map of irrigation areas, Hydrology and Earth System Sciences, 9, 535–547, 2005.

Stocker, B. D., Zscheischler, J., Keenan, T. F., Colin Prentice, I., Peñuelas, J., and Seneviratne, S. I.: Quantifying soil moisture impacts on light use efficiency across biomes, New Phytologist, 218, 1430–1449, 2018.
